# Adaptive Optics in an Oblique Plane Microscope

**DOI:** 10.1101/2024.03.21.586191

**Authors:** Conor McFadden, Zach Marin, Bingying Chen, Stephan Daetwyler, Wang Xiaoding, Divya Rajendran, Kevin M. Dean, Reto Fiolka

## Abstract

Adaptive optics (AO) can restore diffraction limited performance when imaging beyond superficial cell layers *in vivo* and *in vitro*, and as such is of interest for advanced 3D microscopy methods such as light-sheet fluorescence microscopy (LSFM). In a typical LSFM system, the illumination and detection paths are separate and subject to different optical aberrations. To achieve optimal microscope performance, it is necessary to sense and correct these aberrations in both light paths, resulting in a complex microscope system. Here, we show that in an oblique plane microscope (OPM), a type of LSFM with a single primary objective lens, the same deformable mirror can correct both the illumination and fluorescence detection. Besides reducing the complexity, we show that AO in OPM also restores the relative alignment of the light-sheet and focal plane, and that a projection imaging mode can stabilize and improve the wavefront correction in a sensorless AO format. We demonstrate OPM with AO on fluorescent nanospheres and by imaging the vasculature and cancer cells in zebrafish embryos embedded in a glass capillary, restoring diffraction limited resolution and improving the signal strength twofold.

## 1. Introduction

Light-sheet fluorescence microscopy (LSFM) has become a premier tool for volumetric imaging due to its low sample irradiation and high acquisition speed [1]. Its applications include imaging cellular dynamics, organism development and cancer metastasis in 3D model organisms and model systems such as organoids. In a multi-cellular context, optical aberrations are expected to lower the spatial resolving power and sensitivity when imaging beyond one mean scattering free path, which is on the order of ∼100 microns in many tissues. Therefore, LSFM would benefit from adaptive optics [2-4], a technology that was originally developed to compensate atmospheric disturbances for earth bound telescopes but has also found its way into optical microscopy. AO has been successfully applied to raster scanning microscopes [5-7], and has yielded up to an order of magnitude improvement in fluorescence signal in multiphoton systems [8, 9]. It has also found applications in all major super-resolution technologies, namely structured illumination microscopy (SIM) [10-12], stimulated emission depletion (STED) [13, 14] and single molecule localization microscopy [15-20]. However, its adoption in the field of LSFM has been slow. The separate illumination and detection paths in a typical LSFM architecture [1] make the integration of AO complex: both wavefront sensing and correction must be applied individually to the illumination and detection path [21, 22], and additional shift and tilt of the light-sheet plane relative to the detection focal plane need to be corrected separately [21]. While technically possible, the complexity of such a fully AO corrected LSFM system has so far hindered widespread adoption.

Here, we report the application of AO to oblique plane microscopy (OPM) [23], a form of LSFM where the illumination and detection paths share the same primary objective lens. We show that in such a configuration one deformable mirror, conjugate to the pupil of the primary objective, can correct both aberrations affecting the light-sheet and the detection path and assure overlap of the light-sheet plane and the imaging focal plane. This simplifies the implementation of AO in LSFM and dispenses with autofocusing and tilt correction routines proposed for LSFM systems operating in optically complex samples [24]. Further, we introduce “sensorless” AO [25] based on projection images [26], which we found to yield more consistent results than with conventional, static LSFM imaging. The projection imaging removes the bias from finding a suitable image plane to apply iterative optimization on and diminishes the effect of image plane shifts upon modulation of the aberration modes. We demonstrate the working principle of our OPM with AO on test samples consisting of fluorescent nanospheres in agarose and perform AO corrections on the vasculature and cancer cells in zebrafish larvae in a glass capillary in a commercial, automated fluidic screening platform.

## 2. Methods

### 2.1 Adaptive Optics in an Oblique Plane Microscope

Ideally, adaptive optics restores diffraction limited resolution of the imaging system, i.e., it compensates deviations from an ideal spherical wavefront in sample space and deviations from a flat wavefront in Fourier space [4]. Ignoring wavelength dependent effects, it can do so for ingoing (excitation) and outgoing wavefronts (emission) through the principle of time-reversal symmetry and phase conjugation [27]. Therefore, we reasoned that a single corrective element, such as a deformable mirror, could perform AO correction for both the light-sheet excitation and the fluorescence emission in an oblique plane microscope within its shared illumination and detection path.

In Figure 1, a simplified schematic of the primary objective, and an intermediate image plane are shown, representing the front end of an OPM system (the further downstream optics are left out for clarity). In an aberration free scenario (Figure 1A), the oblique light-sheet is launched at the periphery of the objective’s pupil and forms a beam waist which intersects the focal plane. An emitter in the focal plane emits a spherical wavefront, a segment of which (a spherical cap) is converted into a flat wavefront by the objective. Via a tube lens, a diffraction limited image of the emitter is formed. The overall point-spread function (PSF) for the aberration free OPM is the product of the light-sheet intensity distribution and the detection PSF [1], shown in pink at the bottom of Figure 1A. Of note, in an OPM, points that are lying outside of the nominal focal plane and along the light sheet are illuminated and collected too. The objective imprints additional defocus and higher order aberration terms onto their wavefront, which are compensated further downstream via remote focusing [28]. These additional points, and their non-flat wavefronts, are not shown here for clarity.

**Fig. 1.**
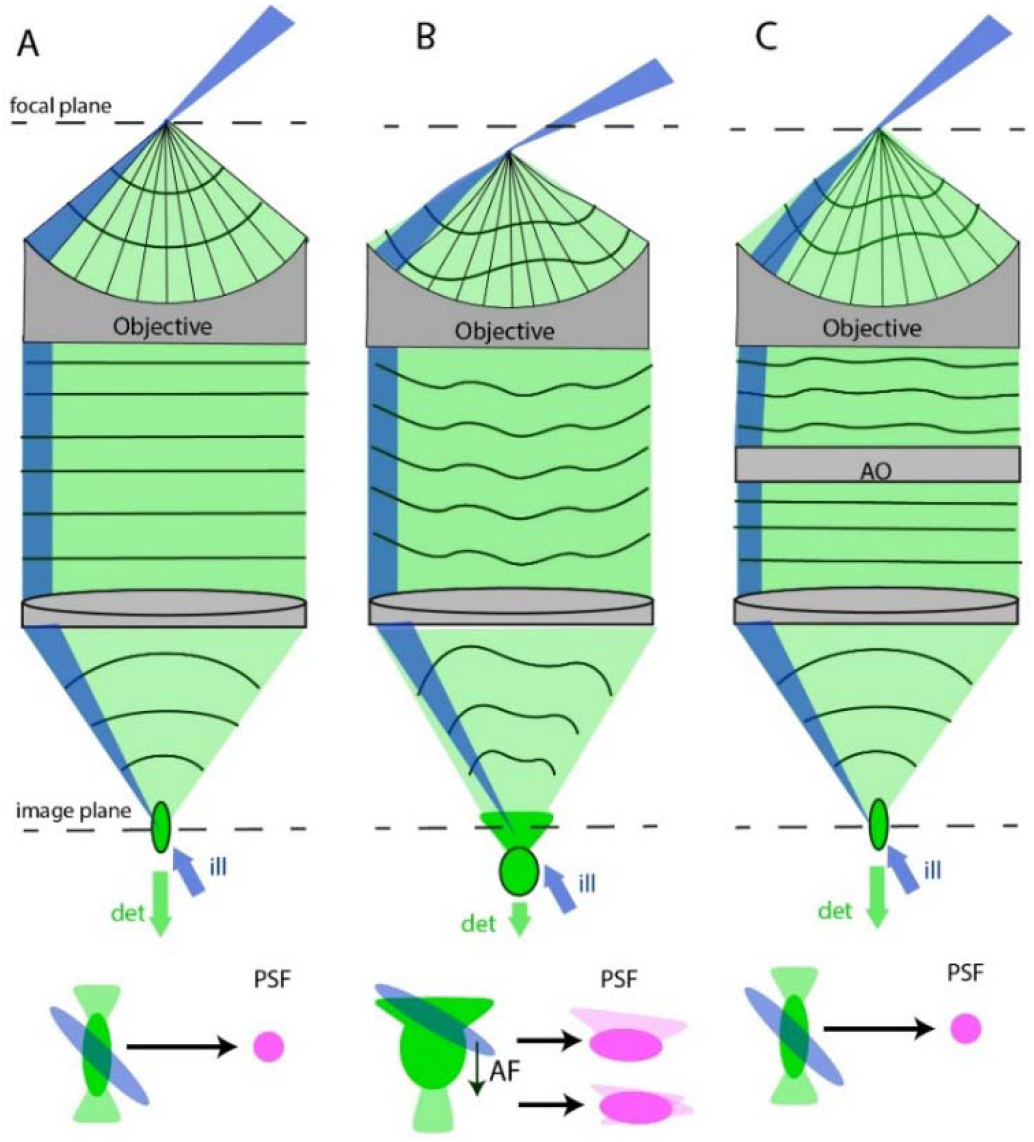
Adaptive optics in an Oblique Plane Microscope (OPM). **A**. Creation of an oblique plane light-sheet (blue) and collection of fluorescence (green) in an aberration free case. The overall point spread function (PSF, pink) is the product of the excitation (blue) and detection PSF (green), shown below. **B**. Imaging in the presence of aberrations: The oblique light-sheet has changed its location and angle due to refraction. Fluorescence emission occurs outside the focal plane, the wavefront is aberrated, and a distorted image is formed outside the image plane. The excitation does not overlap with the best focus of the detection PSF, resulting in a wider overall PSF, shown below. Autofocusing (AF) can re-center the light-sheet but will not undo the overall distortions of the detection PSF. **C**. A corrective optical element, labeled AO, compensates the aberrations of the fluorescence wavefront, restoring a diffraction limited image of the emitter. The same corrective element also imprints a phase correction onto the ingoing light-sheet, such that its distortion in sample space is compensated. The corrective element recovers the diffraction limited PSF of aberration free imaging, shown below.

When aberrations are introduced in sample space, the excitation and fluorescence wavefront is distorted (Figure 1B). As a result, the tilt angle and relative position of the light-sheet to the focal plane is altered. The shown fluorescence emission is distorted by aberrations and originates from outside the focal plane. As such, the image of the emitter is focused outside of the image plane. The overall PSF (shown below in pink) is enlarged, both because the detection PSF is distorted, and because the relative alignment between the light-sheet and the focal plane has been altered. This is a hallmark of spherical aberrations, to give one concrete example, where the marginal rays (to which the light-sheet in OPM belongs) do not overlap with the best focus (which is formed by all rays for the fluorescence detection).

In principle, the light-sheet can be shifted, or in an OPM, as we detail later, one can refocus the detection focal plane separately. This can be done manually or via auto-focusing routines (denoted AF in Figure 1B), but the aberrations imparted on the detection PSF are still present.

Adding a corrective optical element to the beam path corrects the distorted fluorescence wavefront, restoring a diffraction limited detection PSF (Figure 1C). In the other direction, it imprints a phase correction on the light-sheet, which compensates the aberration the sheet experiences in the sample space and thereby re-aligns it with the focal plane. Therefore, the overall OPM PSF is again ideal, i.e. the detection PSF is diffraction limited and intersected at the best focus position by the light-sheet.

This is in contrast with AO in a conventional LSFM geometry, where aberrations for the excitation need to be corrected independently from the detection. This may include an axial shift and a tilt correction to compensate low complexity distortions [24] and higher order aberrations compensation in the detection path [21]. Thus, OPM combined with AO has the potential to reduce the number of optimization (or wavefront sensing) steps, which can save time and fluorescence photons, and to reduce the overall complexity of the setup.

### 2.2 Optical setup

In a typical implementation of adaptive optics, a corrective element, such as a deformable mirror, is conjugated to the pupil of the objective [4]. This is also the place where a galvanometric mirror is placed to allow rapid volumetric scanning in OPM [29, 30]. While additional relay optics could conjugate both a deformable and galvanometric mirror to the same pupil, this would increase the number of optical elements, which in turn would make the system more complex to build and increase light-losses by each additional lens. Instead, we used a galvo pair in an image plane to perform volumetric scanning [31], and placed the deformable mirror in the Fourier space. As such, the number of lenses is the same as in a traditional OPM implementation.

A schematic overview of the system is shown in Figure 2. Excitation light, shown in blue, from CW lasers (LightHUB Ultra light engine, equipped with 488nm: 200mW, 561nm: 150mW, 642nm: 140mW, Omicron-Laserage Laserprodukte, Germany) is delivered via a single mode fiber to an illumination engine, which was introduced by Chen *et al*. [32]. After collimation, the beam passes through a Powell lens (10□ fan angle, Laserline Optics, Canada) and is collimated in one dimension by an achromatic doublet (f=30mm, Thorlabs, L1 in Figure 2A), and focused into a light-sheet in the other dimension. A relay lens pair (L2 and L3) images the light-sheet onto a resonant galvo (CRS4K, Novanta). After passing through an achromat lens (L4, f=50mm, Thorlabs), the laser light is coupled into the main optical train of the OPM via a dichroic mirror (Di03-R405/488/561/635-t1-25×36, Semrock). The excitation light passes over a deformable mirror (Mirao52, Imagine Optic), which is conjugate to the pupil of the primary objective (O1 in Figure 2A: NA 1.10, Apo LWD 25X, Water-dipping, MRD77220, Nikon). The laser beam is scanned by two galvo mirrors (Galvo 1 and 2, GVS011, Thorlabs) placed in an image space, before being launched by O1 at an oblique angle.

**Fig. 2.**
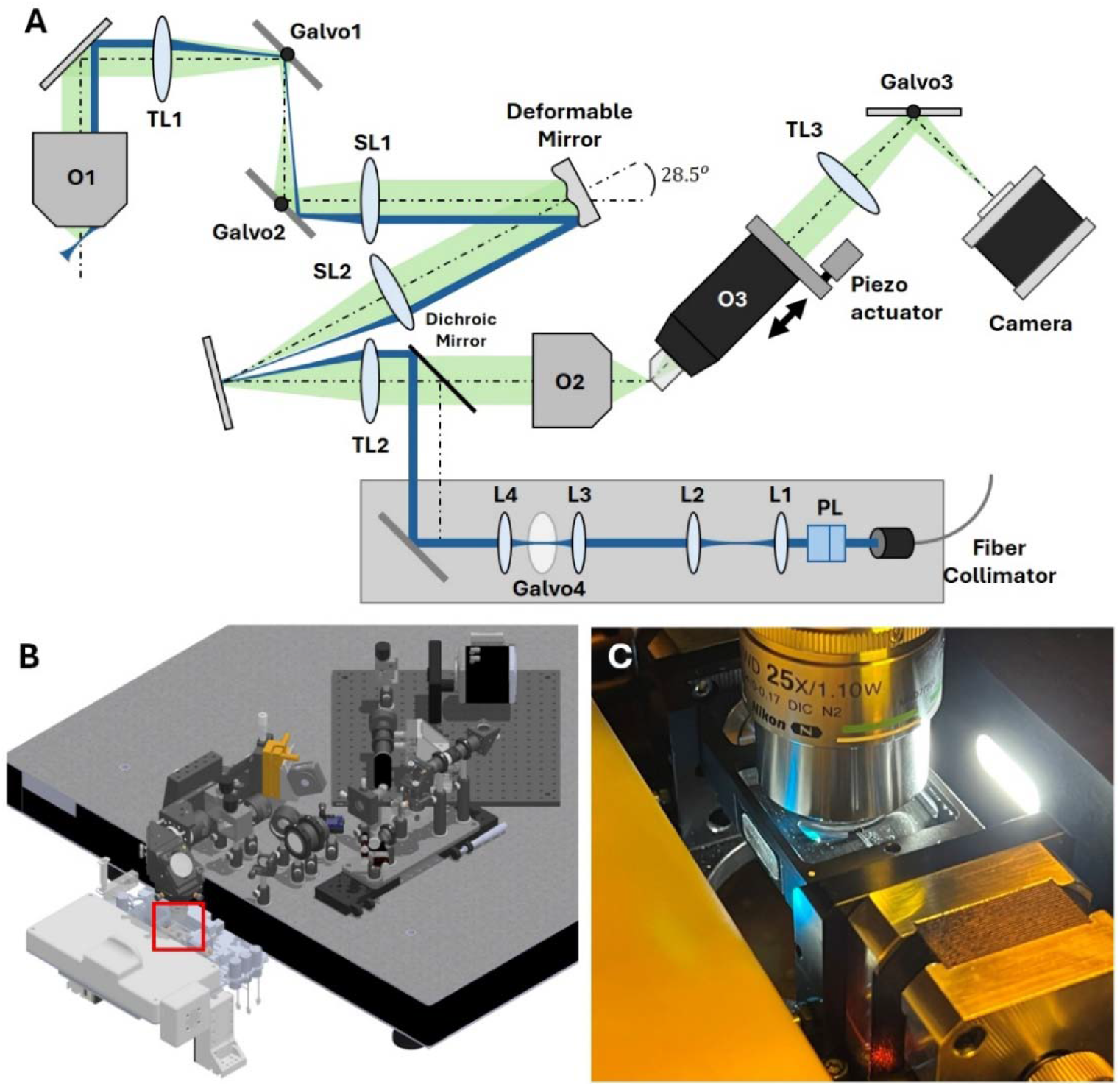
Setup for Oblique Plane Microscopy with adaptive optics. **A**. Schematic overview of the setup. **B**. Rendering of a CAD model of the setup. The optical microscope is assembled on an elevated breadboard and the VAST BioImager system is shown in white and light gray to the left. **C**. Photograph of the experimental setup, corresponding to the boxed red region in B. PL: Powell lens, TL: Tube lens,SL: Scan lens, L: Achromatic doublet lens, O: Objective.

Fluorescence light (shown in green) is collected by the primary objective and after passing through a tube lens (TL1, TTL200MP, Thorlabs), it is de-scanned by the two galvo mirrors. After passing through a scan lens (SL1, f=165mm, TTL165-A, Thorlabs), the fluorescence light is reflected by the deformable mirror. The second scan lens (SL2, identical to SL1) and second tube lens (TL2, f=169 mm lens assembly, consisting of TTL200MP and AC508-750-A, Thorlabs) map the deformable mirror to the pupil of the secondary objective (O2: UPlanXApo, 20X, NA 0.8, Air, Olympus). O1 and O2 form a remote focusing system [28], and as such, the fluorescence light is focused by the secondary objective into a distortion free 3D image of the sample space. A tertiary imaging system, rotated by 45° and consisting of a glass-tipped third objective (O3: AMS-AGY v2, NA 1.0, Applied Scientific Instrumentation), a tube lens (TL3, f=300mm, ACT508-300-A-ML, Thorlabs) and a galvo mirror (Galvo3, GVS 011, Thorlabs), images the fluorescence light onto a camera (Orca Flash 4.0, Hamamatsu). The tertiary objective is actuated by a piezo actuator (POLARIS-P20A, Thorlabs) with 17 microns range, both to compensate for thermal drift and to refocus the detection path separately from the illumination path.

The microscope system is built on an elevated platform, with a CAD rendering of the setup being shown in Figure 2B. The primary objective interfaces from the top with the sample chamber of a vertebrate automated screening technology (VAST Bioimager, Union Biometrica) platform (Figure 2C). The VAST system allows automated loading and positioning (XY axes and longitudinal rotation angle) of 2-7 days post-fertilization (dpf) zebrafish larvae. A glass capillary (700-micron diameter, 40-micron wall thickness) ensures mechanical stability, but also introduces aberrations that impair high-resolution imaging. While the system is designed for screening of zebrafish larvae combined with high-resolution imaging of regions of interest, the OPM system in an upright configuration could conceivably be used for other applications, such as intravital imaging of superficial layers of the mouse cortex.

Microscope control is performed via an open source Python-based software package called navigate [33], where the deformable mirror control and projection imaging was implemented as a plugin feature. The navigate software also has a generic autofocusing routine, which we used on our OPM using the piezo actuator on the tertiary objective. Triggering and waveform control of the camera and galvanometric mirrors is performed via a NI Data Acquisition card (PCIe-6738, National Instruments). The deformable mirror was calibrated using a Shack-Hartmann wavefront sensor (Haso4, Imagine Optic) and an interaction matrix was established using WaveTune (Imagine Optic). Mirror control was then implemented via navigate using the WaveKit SDK from Imagine Optic, and we used a decomposition of the wavefront into 32 Zernike modes. Throughout this work, we used the Shannon entropy of the normalized discrete cosine transform (DCTS) [24] as the image metric to optimize the Zernike modes, and for autofocusing.

### 2.3 Projection imaging for sensor less adaptive optics

Our adaptive optics systems in this work dispensed with a wavefront sensor, but instead employed an iterative optimization scheme for wavefront modes, commonly referred to as sensorless AO [25]. In such a scheme, the wavefront is decomposed into a finite, orthogonal set of modes (typically Zernike modes), and the coefficients for each mode are optimized one by one using an image metric, i.e., for each coefficient, a series of 2D images is acquired for different values of the coefficient, and an image metric is computed on the fly.

In this work, each Zernike coefficient was probed with 3-5 values, and a parabolic fit of the corresponding image metric approximated the optimal value of the coefficient. The corresponding correction was added to the wavefront on the mirror prior to optimizing the next mode. In this way, each successful correction of a mode increased the signal strength for the subsequent mode optimization [34]. Once the full set of modes (typically 29, where we left out the first three Zernike modes, “tip”, “tilt”, “defocus”) are optimized, termed an iteration in this work, the scheme may be iterated again to improve the convergence of the wavefront. The data presented was obtained by performing 3 iterations in total.

Sensorless AO involves a user bias to select an appropriate image plane which contains features that are responsive to the image metric. For example, for an image sharpness or intensity metric, the selected image plane should contain sharp or bright features, respectively. If it does not, the AO iteration will slowly (or not at all) converge due to noisy images and correspondingly unreliable image metrics. In addition, Zernike modes and imperfections of the mirror to reproduce them can lead to a loss in orthogonality and unwanted modal crosstalk [35]. Therefore, the image plane may get axially and laterally displaced during the optimization routine, even though we exclude these lower modes (tip/tilt/defocus) from our iteration. In an in-homogeneously labeled sample, this can lead to additional “noise” in the optimization, as the algorithm may want to move the focal plane to sharper or brighter features outside of the selected image plane.

To counter the selection bias and focal plane jitter, we hypothesized that a recently introduced optical implementation of the shear warp transform [36], which can produce projection images from different viewing angles in a single camera exposure, could be used for sensorless AO optimization. Figure 3 schematically shows the working principle of the projection method. In Figure 3A, the primary objective is shown, with the light-sheet and the detection focal plane being tilted to the optical axis (denoted as z). Using the galvo scanning unit, the light-sheet and the focal plane can rapidly sweep the sample, covering the parallelepiped-shaped volume shown in light blue. When that sweep is completed during one camera exposure, a sum projection of this volume is formed (Figure 3B).

**Fig. 3.**
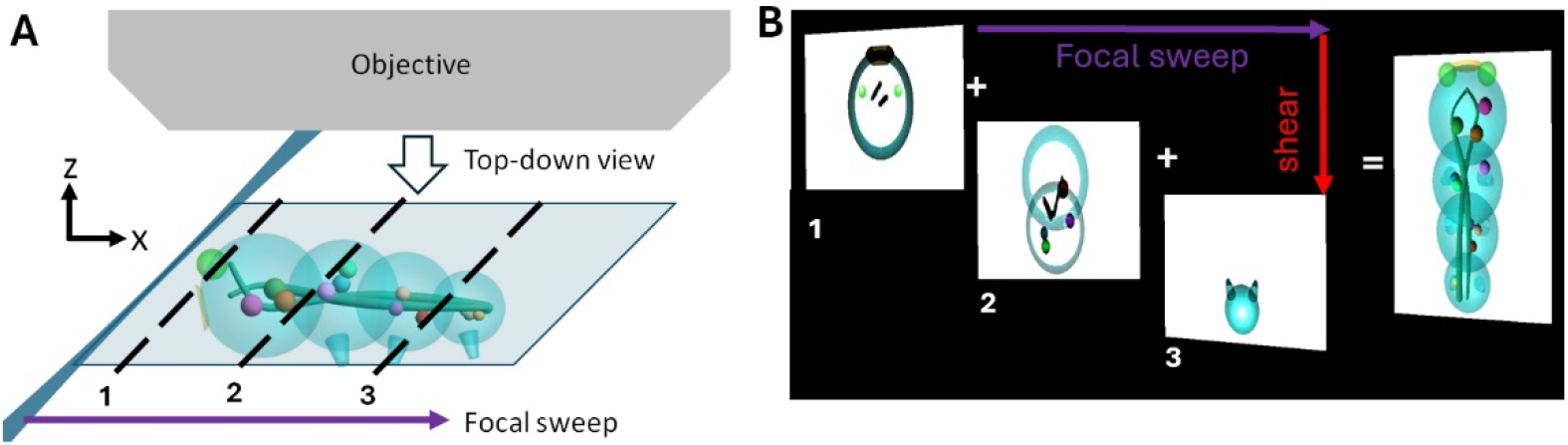
Illustration of the multi-angle projection method. **A**. Schematic illustration of a sample, the primary objective and a light-sheet (shown in blue). A focal sweep (violet arrow) is obtained by rapidly scanning the light-sheet and the corresponding focal plane so that they span the light-blue shaded volume. The dashed lines show three individual positions, labeled 1-3, along the focal sweep. **B**. When the focal sweep occurs during one camera exposure, a projection is formed (right). Adding additional shear (red arrow) during the focal sweep changes the viewing direction of the projection. Panels 1-3 show instantaneous images along the focal sweep, corresponding to the dashed lines in **A**. The added shear in this illustration results in a top-down view, with the viewing direction indicated with an arrow in **A**

Figure 3B shows the addition of image shearing introduced during a focal sweep in more detail. As the light-sheet is scanned through the sample, the immediate (instantaneous) images (three representative image planes are shown along the dotted lines in Figure 3A) are shifted over the camera sensor during one exposure. Per the shear warp transform, a projection along a different viewing direction can be formed. In the example of Figure 3B, a “top-down” view (arrow in Figure 3A) is formed, but in principle other viewing directions are possible, depending on the amount of image shear that is applied. Physically, the amount of shear is controlled by the scan amplitude of the so-called shear galvo (Galvo3, Figure 2A), whose frequency is matched to that of the focal sweep.

In this manuscript, we use the term “static AO” when images are acquired with stationary scanning mirrors during an iterative wavefront optimization, and “projection AO” when a focal sweep and shearing is performed, respectively. A top-down view of a 38 µm sweep was used for all datasets shown, although projections of arbitrary viewing angles and other projection depths could be used as well.

## 3. Results

### 3.1 AO correction of fluorescent nanospheres in a glass capillary

To evaluate the effectiveness of the AO correction in an OPM, we imaged fluorescent nanospheres (100 nm diameter, yellow-green FluoSpheres carboxylate, Invitrogen) diluted 1:2000 in 2% agarose in a glass capillary (40-micron wall thickness, 700-micron diameter) of the VAST bioimager system. We first applied the system correction to the deformable mirror, which is an AO correction that we optimized on fluorescent nanospheres adherent on a glass coverslip, i.e. an aberration free sample. Similarly, the relative alignment between light-sheet and focal plane was also optimized beforehand on this coverslip sample.

With the system correction applied, the tertiary objective had to be re-focused by about 3 microns when imaging in the glass capillary, which corresponds to a sample space mismatch of ∼2.3 microns (the sample space is demagnified uniformly by a factor of 1.333 compared to the remote space). Even after this optimization, strong aberrations are visible in the nanosphere data. Figure 4A shows maximum intensity projections (MIP) of a stack that was acquired after the autofocus optimization. We estimated the resolution to 520nm±49nm (n=21) using image decorrelation analysis [37]. This value was below the performance of such an OPM, which has previously been reported to achieve sub-400nm resolution [32].

**Fig. 4.**
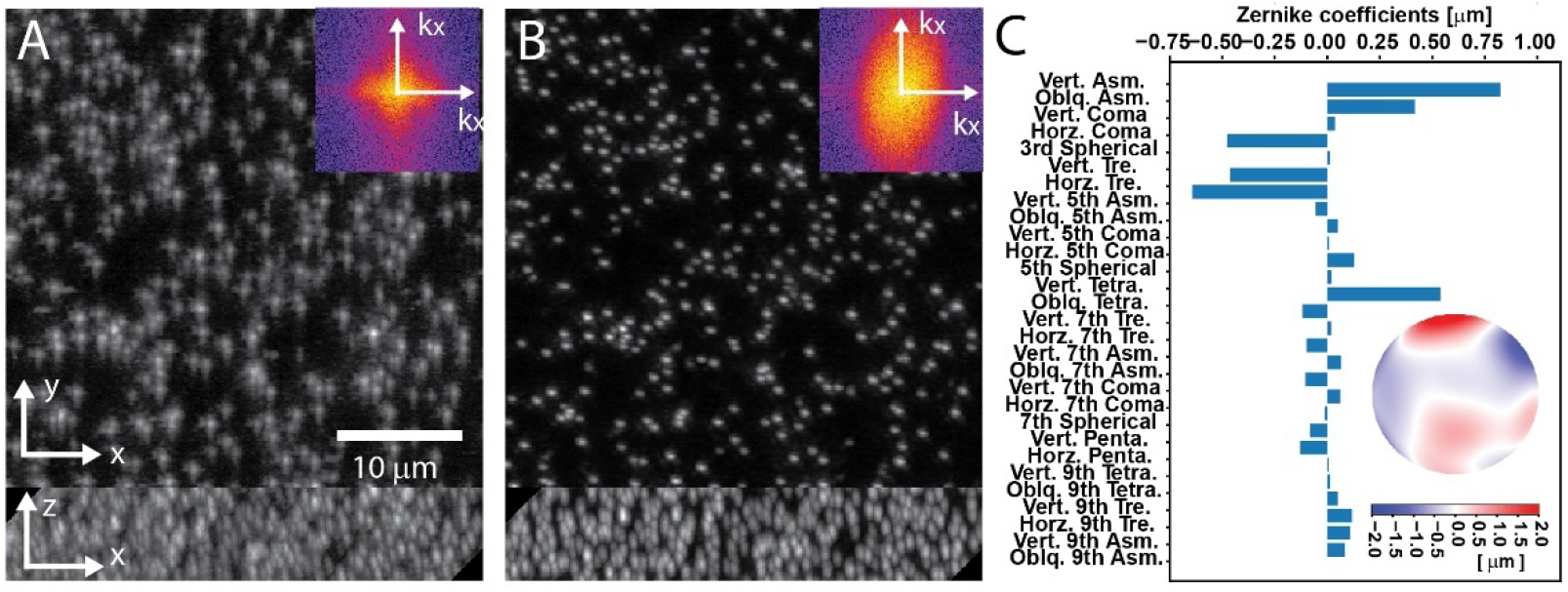
Adaptive optics on fluorescent nanospheres embedded in a glass capillary with agarose. Maximum intensity projections of 3D stacks of nanospheres are shown **A**. without and **B**. with AO-correction. Inlays: logarithm of the power spectral density of the x-y images. **C**. Zernike coefficients for the measured AO-correction, along with the wavefront (inlay). Zernike coefficient abbreviations: ASM: Astigmatism, TRE: Trefoil, Tetra: Tetrafoil, Penta: Pentafoil, Vert: Vertical, Oblq: Oblique.

When we applied the sensorless correction scheme, using static AO, the beads in the images became much more compact, and the peak signal increased approximately twofold (Figure 4B). Similarly, the power spectrum had a larger extent, and we estimated the lateral resolution to 376±23nm (measured over 21 planes of each stack) using image decorrelation analysis. Notably, the power spectrum after AO correction has an anisotropic support, which is typical in OPM systems as the resolution in the tilt direction of the tertiary objective is lower [38].

Importantly, we started the AO correction without autofocusing of the light-sheet to test if the wavefront correction would also fix the relative alignment between the focal plane and the sheet. After the AO optimization was completed, we ran the autofocusing routine. Less than half a micron of re-alignment was needed (∼0.38 microns in sample space), which is well within the depth of focus of the detection system.

### 3.2 Comparison of sensorless AO using static and projection imaging

Next, we compared static AO optimization to AO optimization with projection imaging. To this end, we imaged the same fluorescent nanosphere sample at 20 different locations along the central axis of the capillary. Positions were spaced apart by 400 µm, large enough to prevent overlap of scanned areas. Figure 5A shows MIPs of 3D stacks at one position using system, static and projection AO corrections. Notably, the individual Zernike coefficients showed more variability across all positions with the static AO compared to projection AO (Figure 5B).

**Fig. 5.**
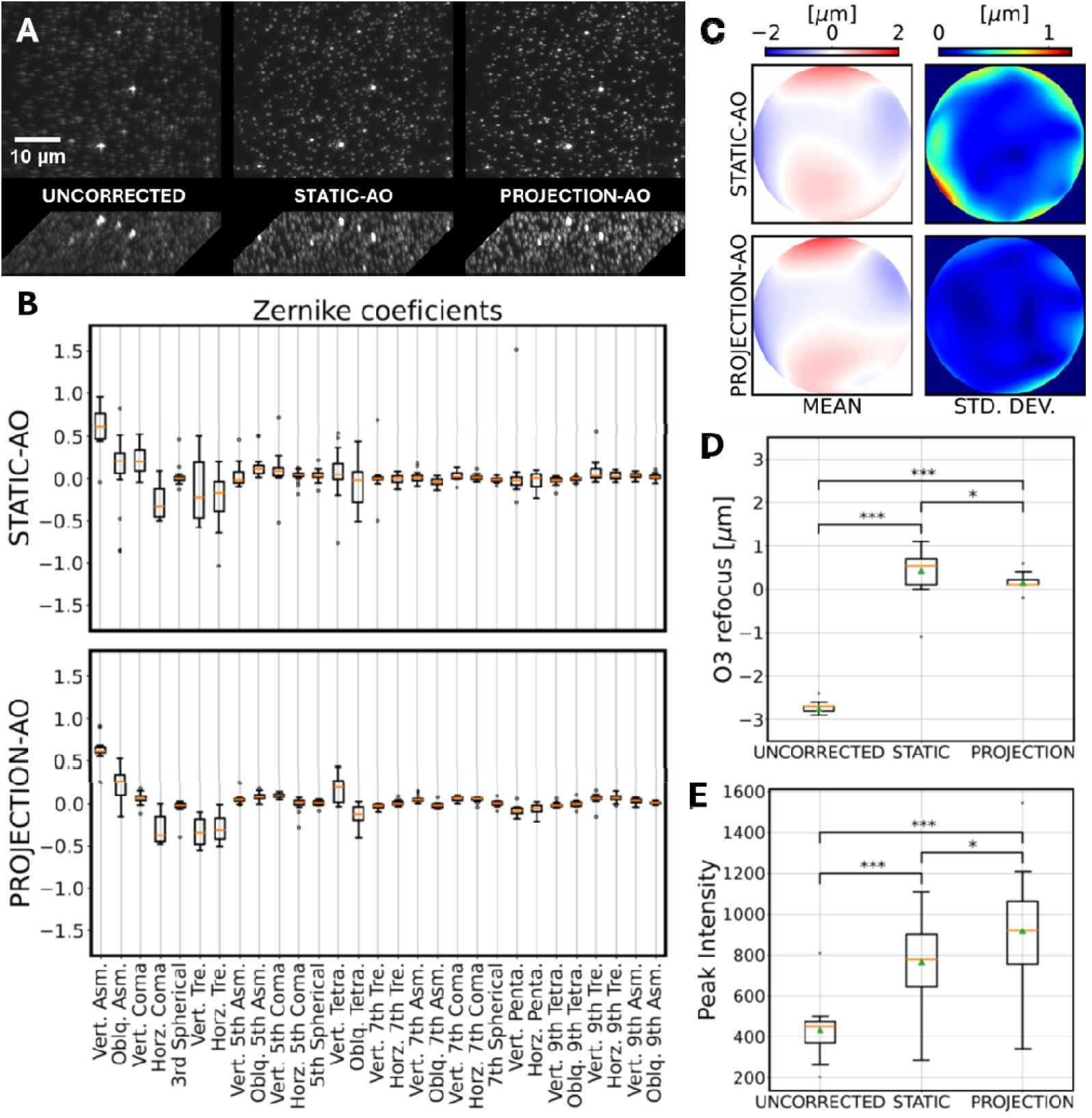
Performance of iterative adaptive optics with static and projection imaging using fluorescent nanospheres embedded with Agarose in a glass capillary. **A**. Fluorescent nanospheres, as imaged with system, static AO, and projection AO correction, respectively. Maximum intensity projections of 3D stacks are shown. **B**. Overall Zernike coefficient amplitudes for 20 different measurements along the capillary length using static and projection AO, displayed as box plots. **C**. Mean (left) and standard deviations (right) of all wavefronts measured using (top) static AO and (bottom) projection AO. **D**. Focus correction required for either the uncorrected case or post-AO correction to reach the highest DCTS image quality. **E** Peak bead intensity for all measurements and AO conditions. Zernike coefficient abbreviations: ASM: Astigmatism, TRE: Trefoil, Tetra: Tetrafoil, Penta: Pentafoil. Vert: Vertical, Oblq: Oblique. For statistical testing, alternate hypotheses were (for example) **D:** H_1_ = |Proj| < |Stat| and **E:** H_1_ = Proj > Stat. A one-sided Mann-Whitney U test was used for all calculations. Significance is notated as *: p < 0.05, **: p < 0.01, ***: p < 0.001. Mean values are represented by green triangles in **D-E**

Moreover, while both static and projection imaging produced a similar mean wavefront correction, the standard deviation was considerably higher in the static modality compared to the projection imaging (Figure 5C). Assuming that the aberrations introduced by the glass capillary are stereotypical and that fluorescent nanospheres in Agarose make for a reasonably homogeneous sample, we expected a similar wavefront correction regardless of the measured position. Consequently, we assume that projection AO provided a more accurate, less biased picture of the aberrations at a given location. In addition, the more consistent wavefront correction also resulted in a tighter autofocus correction value overall, an effect that was statistically significant, as shown in Figure 5D. The focus correction for projection AO was low enough in both its variance and magnitude that it effectively makes the autofocus routine redundant.

Finally, since the Strehl ratio is related to the root-mean-square deviation of the wavefront [39, 40], a better AO correction is expected to lead to a higher peak intensity in the nanosphere image. For a given position/condition, peak intensity was measured from the images as the median bead intensity for the top-50 brightest beads (Figure 5E). While both methods yielded a marked improvement over system correction with autofocus alone, projection AO notably achieved a significantly higher peak intensity than static AO.

We performed similar measurements over 33 hand-chosen positions by imaging Zebrafish vasculature, and performance was evaluated between the two AO modes (Supplementary Figures 1-3). The Zebrafish vasculature was labeled with *Tg(kdrl:EGFP)* in a casper background. Zebrafish husbandry and experiments followed established protocols and have been approved and conducted under the oversight of the Institutional Animal Care and Use Committee (IACUC) at UT Southwestern under protocol number 101805. Fish were sacrificed and fixed in 4% paraformaldehyde (PFA) at 5 days post-fertilization (dpf) and suspended in the VAST capillary in 0.2-0.5% low-melting point (LMP) agarose for imaging. Various regions were imaged throughout the fish, although most were in the tail.

Like the bead measurements, we found that the variation in the Zernike coefficients appeared lower in Projection AO, and the standard deviation in the wavefront was lower as well (Supplementary Figure 1). Notably, for some locations, the region of interest remained steadier during AO iteration for projection AO compared to static AO (Supplementary Figure 2 and Supplementary Movie 1). We assume that modal coupling of higher aberration modes, and imperfections of the deformable mirror, leads to defocus and lateral shifting, which manifests itself in image jitter in Supplementary Movie 1 for static AO. In contrast, in the projection mode, features remain within the region of interest, and they get progressively sharper. The result of this jitter effect can be seen in Supplementary Figure 3B, where static AO failed to converge to a good wavefront correction.

Figure 6 shows the result for one of the 33 Zebrafish imaging experiments. In Figure 6A, MIPs of a 3D stack that was acquired with the system correction applied to the mirror, and after running the autofocusing routine. The image appeared unsharp, and many details of the vasculature were blurred out. Figure 6B shows MIPs of a stack acquired after three rounds of projection AO optimization. Much finer details of the vasculature were visible, including clearly delineated nuclei of the endothelial cells. Figure 6C shows the corresponding wavefront correction (i.e. after subtracting the system correction) and the Zernike coefficients below. Supplementary Figure 3A shows all 33 individual wavefronts.

**Fig. 6.**
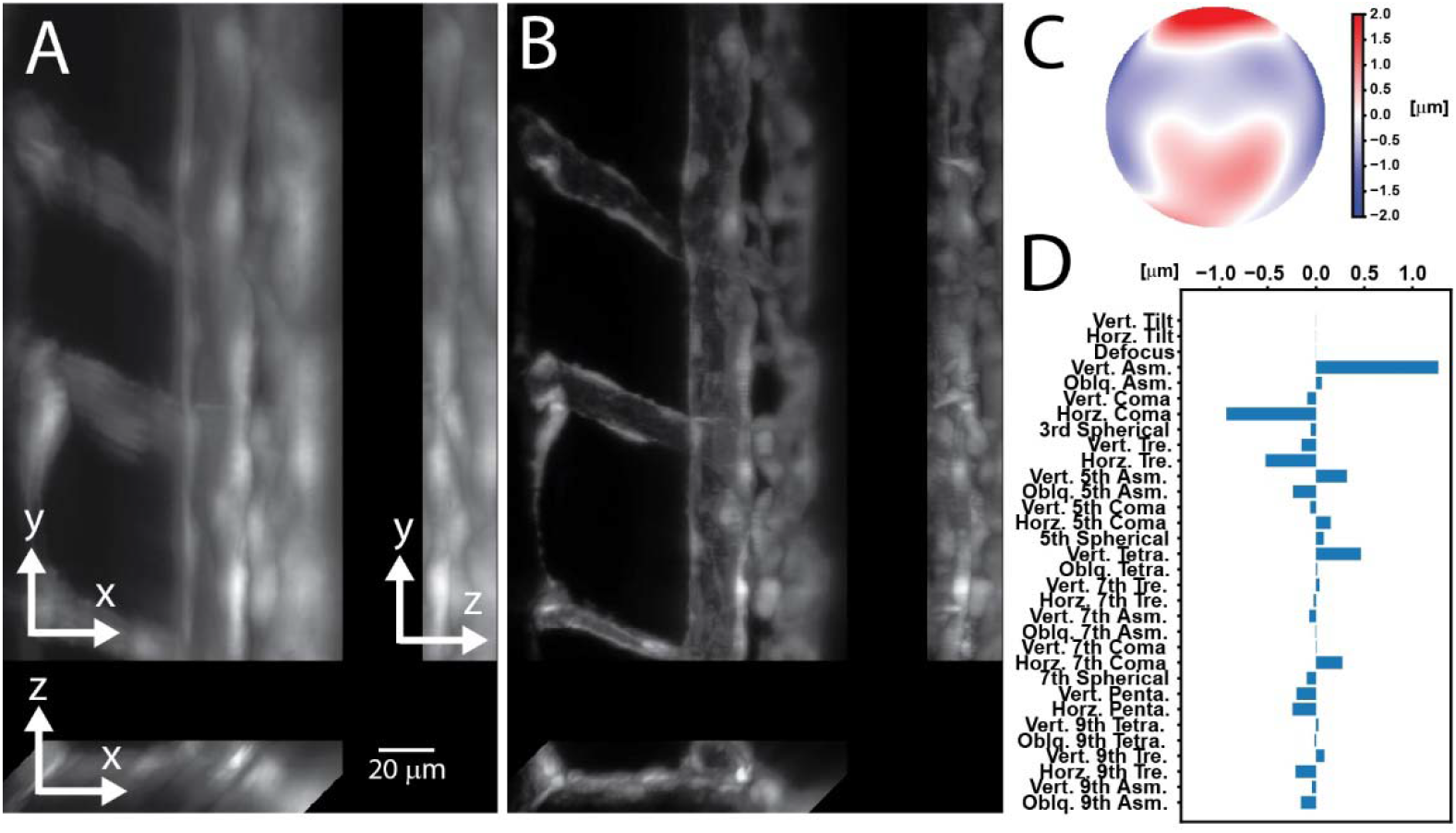
Projection based adaptive optics in a zebrafish larva, labelled with the vasculature marker Tg(*kdrl:EGFP*). **A**. Maximum intensity projections (MIPs) of a stack acquired with the system correction applied to the mirror, and after performing the autofocusing routine. **B**. MIPs of a stack acquired after projection AO correction. **C**. The wavefront correction applied in **B D**. List of the corresponding Zernike coefficients. Zernike coefficient abbreviations: ASM: Astigmatism, TRE: Trefoil, Tetra: Tetrafoil, Penta: Pentafoil.Vert: Vertical, Oblq: Oblique.

While the glass capillary is necessary for the operation of our VAST system, many zebrafish LSFM imaging experiments are conducted in fluorinated ethylene propylene (FEP) tubes, which have a refractive index close to water. In Supplementary Figure 4, we compare the performance of static and projection imaging when imaging live Zebrafish in an FEP tube. 3 dpf fish were suspended alive in the FEP tube in 0.2% LMP agarose and 0.01% Tricaine as anesthetic. 15 positions were measured, imaging the vasculature. While the improvements were less substantial than in the glass capillary, there was still a notable increase in image sharpness relative to system correction with autofocus alone. Notably, both static and projection AO were successful in correcting the images and refocusing light sheet, despite the presence of motion artifacts from blood flow.

### 3.3 Zebrafish Xenograft imaging

To showcase the potential of adaptive optics with OPM for subcellular imaging of cancer cells in a zebrafish xenograft, we imaged a zebrafish larva injected with A375 human melanoma cells expressing the actin marker pLVX-iNeo-mRuby-Tractin. For tissue context, the zebrafish expressed the vascular marker Tg(*kdrl:eGFP*). A375 cancer cells were injected at 2 dpf into the yolk near the common cardinal vein (CCV). Cancer cells were allowed to metastasize for 3 days and then the zebrafish were sacrificed and fixed at 5 dpf and mounted in 0.5% LMP agarose in a 700 µm diameter glass capillary. AO correction was performed in the projection mode with a 38 µm sweep using the cancer cells in the red fluorescence channel as the target.

Figure 7 shows MIPs of stacks acquired in the caudal hematopoietic tissue (CHT) region before and after the application of AO, highlighting the improvements for imaging zebrafish vasculature and A375 cells. Importantly, fine details were recovered in the vasculature in the green channel (Figure 7C and Movie 2) despite sensorless AO being performed only on the cancer cells in red channel (Figure 7D).

**Fig. 7.**
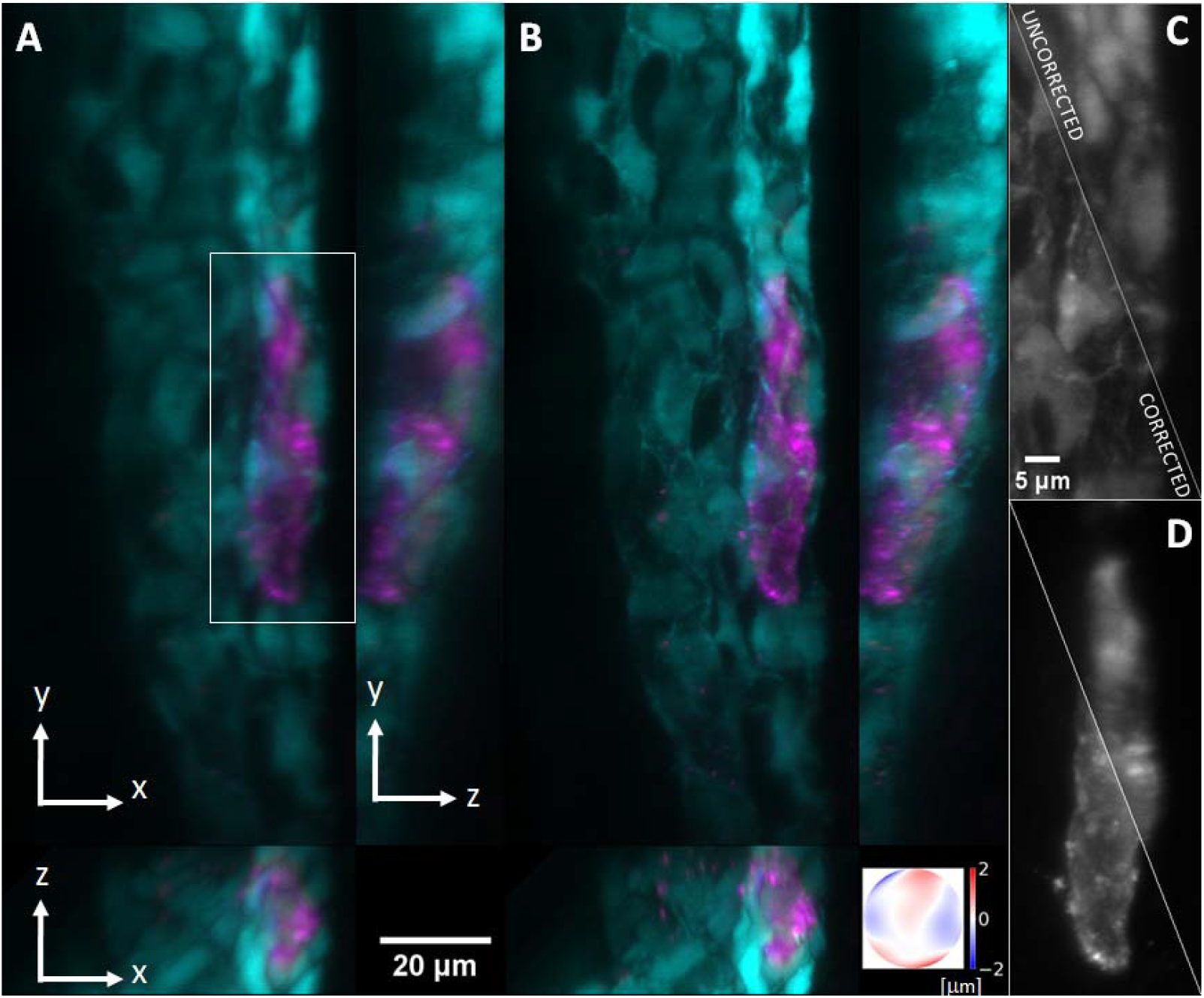
Projection AO for subcellular imaging of cancer metastasis in zebrafish xenografts. The xenografted zebrafish was mounted in agarose in a glass capillary, and the CHT region was imaged. **A**. Uncorrected XY, XZ and YZ MIPs of vasculature Tg(kdrl:eGFP) (cyan) and A375 cancer cells labelled with the actin marker pLVX-iNeo-mRuby-Tractin (magenta**). B**. Projection AO-corrected stacks, with the measured wavefront show in the inlay (color bar is the wavefront error in µm). **C**. Zoomed-in regions (ROI designated in A) for the vasculature and **D**. cancer cells emphasize the improvement in fine details after AO correction. Note that D roughly shows the ROI on which projection-AO was performed.

## 4. Discussion

In this work, we have introduced static and projection-based adaptive optics for oblique plane microscopy (OPM), where one deformable mirror corrects both the light-sheet and the fluorescence detection path. This reduces the complexity for AO integration into a light-sheet microscope compared to previous implementations, which have involved separate AO systems for the excitation and detection paths. We demonstrate practical applications for imaging of zebrafish larva in a commercial fluidic screening platform. The aberrations introduced by the glass tube are stereotypical, which allowed us to carefully compare the convergence of the AO iterations using both static and projection imaging on test samples and Zebrafish embryos. The imaging results and statistical analysis support the assumptions taken for our system, namely that dispersion and wavelength effects are low, and the AO correction can be applied to volumetric imaging of the OPM.

We show that a successful AO correction in our OPM system overlaps the light-sheet with the detection focal plane. This was expected from an ideal wavefront correction (i.e., that it works in both directions), and we have shown that the effect is repeatable. This contrasts with a traditional light-sheet system with separate illumination and detection lenses. Upon adaptive correction, the light-sheet and the focal plane of the detection system are not guaranteed to overlap, as both the sample, as well as the relative AO correction may introduce lateral and axial shifts. As such, previous LSFM AO systems still needed a separate “auto-focus” routine to overlap the sheet with the focal plane. In our case, such a routine is no longer necessary, and as such makes the system more robust and potentially faster.

With OPM’s optical scan mechanism to rapidly scan a 3D sample, it is compatible with multi-angle projection imaging at camera limited rates. As such, we explored if projection imaging could be beneficially applied in an iterative, sensorless AO modality where an image metric is optimized. We found that projection AO enables more consistent wavefront estimation, and that the effect is statistically significant. We attribute this effect to less fluctuations caused by mode coupling and related re-focusing and shifting of a single image plane. Furthermore, projections reduce the bias on selecting the right image plane to perform AO on, making it a more robust tool for fast corrections.

We performed “top-down” projections for sensorless AO optimization. They provide the overall sharpest view as the projection axis corresponds to the most elongated axis of the detection PSF (the z-axis in Figure 3A). We assumed that the DCTS metric would then also be the most sensitive. Nevertheless, one could also use different viewing directions, potentially optimized for each wavefront mode. As a naive example, spherical aberrations break the axial symmetry of the PSF, and as such a “side-view” may contain more useful information than a top-down view. This could also be leveraged with machine learning algorithms to estimate aberrations.

We applied sensorless adaptive optics for simplicity, but OPM AO could also be implemented with guide star-based [41-43] or other direct wavefront sensing strategies [44]. Compared to traditional light-sheet architectures, our approach would still introduce the advantage of requiring only one guide star and wavefront sensor. Besides speed gains (i.e. direct wavefront sensing can in principle determine a correction with one camera acquisition), potentially the full pupil of O1 can be observed, and corrected. In the current implementation, the detection path clips some portions of O1 pupil due to the tilting of the tertiary arm. We assume that due to the smoothness of the lower order Zernike modes, the correction we iteratively determine still corrects well for the portions that are not visible to the detection system.

Importantly, all data shown within this manuscript are raw data, unlike previous lattice light-sheet work where AO was combined with iterative deconvolution, such as the Richardson Lucy (RL) method [21]. We opted not to apply iterative deconvolution methods here to let the reader better distinguish the gains obtained from adaptive optics alone. Nevertheless, based on previous deconvolution results on a similar OPM platform, it is expected that the resolution could be improved to ∼300 nm scale with for example RL type deconvolution [32].

In summary, we found that an OPM system, as it uses only one primary objective for illumination and fluorescence detection, lends itself to adaptive optics. We managed to successfully restore high-resolution imaging in a commercial fluidic zebrafish screening platform and have also shown improvements in a more common FEP tube mounting system. Given the vast and yet unexplored engineering landscape offered by combining AO and OPM, it is anticipated that significant further advancements await.

## Acknowledgments

The authors are grateful to Dr. Vasanth Siruvallur Murali for help with zebrafish xenografting. The authors are also grateful to the Danuser lab at UT Southwestern for their help on computational support and the Animal Resource Center (ARC) at UT Southwestern for zebrafish husbandry and maintenance. Moreover, this research was supported in part by the computational resources provided by the BioHPC supercomputing facility located at UT Southwestern Medical Center.

## Disclosures

The authors declare no conflict of interest.

### Data availability

Data underlying are available upon request.

### Supplemental document

See Supplement 1 for supporting content.

