## Supplement 1 for "Adaptive Optics in an Oblique Plane Microscope"

Adaptive Optics in Oblique Plane Microscopy: supplemental document

**Supplementary Figures**

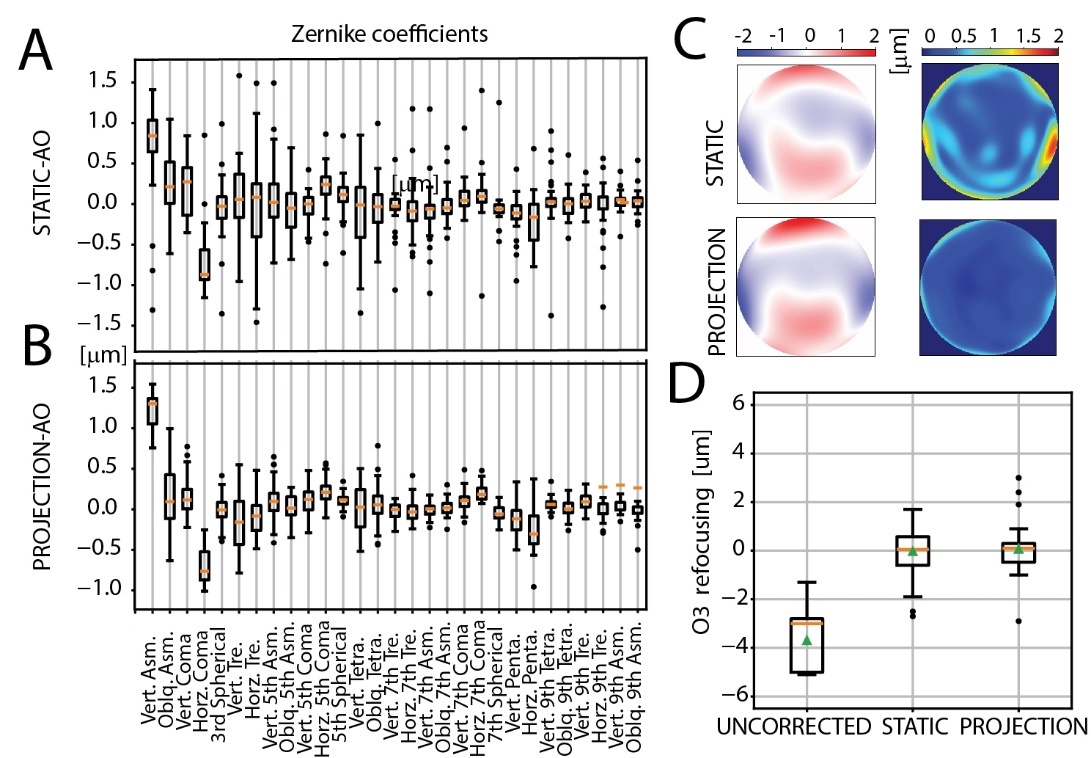

**Supplementary Fig. 1.** Performance of iterative adaptive optics evaluated in imaging larval zebrafish with labelled vasculature mounted in a glass capillary, comparing static light-sheet and projection imaging. Corrections were performed for 33 positions in the capillary. **A. and B**. Overall Zernike coefficient amplitudes for all positions measured using the. static and projection modes are displayed as box plots. **C**. Mean (left) and standard deviations (right) of the wavefronts measured using (top) static-AO and (bottom) projection-AO methods **D.** Focus correction required for either the uncorrected case or post-AO correction to reach the highest DCTS image quality. Zernike coefficient abbreviations: ASM: Astigmatism, TRE: Trefoil, Tetra: Tetrafoil, Penta: Pentafoil. Vert: Vertical, Oblq: Oblique.

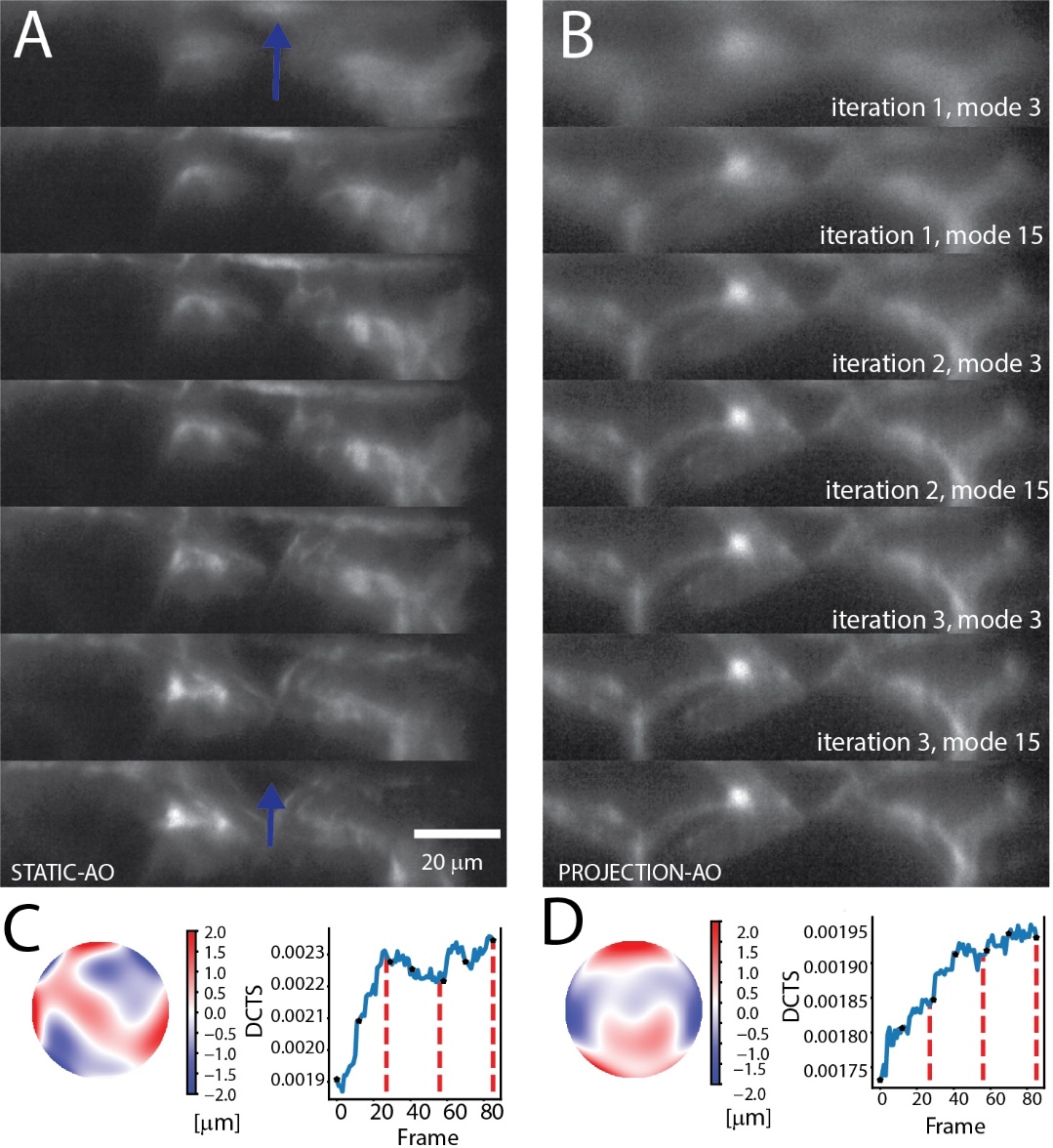

**Supplementary Fig. 2. A. and B**. Individual frames from the AO routine for one Zebrafish vasculature imaging location (Position 6) for **A.** static and **B**. projection modes. **C and D** Relative increases in the DCTS metric are shown below the frames, with black dots indicating the displayed frames and red lines marking the end of each iteration, along with AO-measured wavefronts.

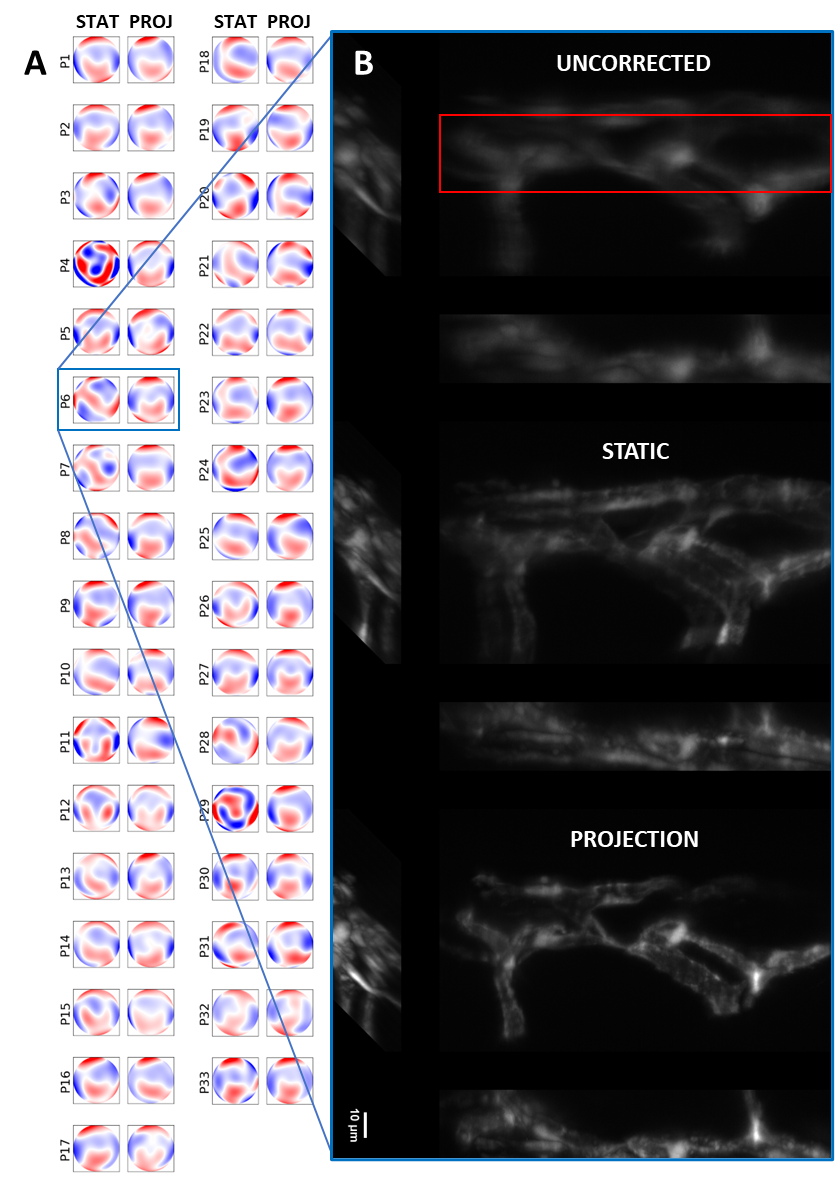

**Supplementary Fig. 3.** More detailed comparison of static vs projection AO on fixed Zebrafish embedded in a glass capillary. **A.** Measured wavefronts for all positions (labelled P1-P33) for both static and projection modes. **B.** 3D stack MIPs for the case of P6 where static AO failed. The ROI over which the sensorless AO was run is marked in red and is the same ROI shown in Supplementary Figure 2.

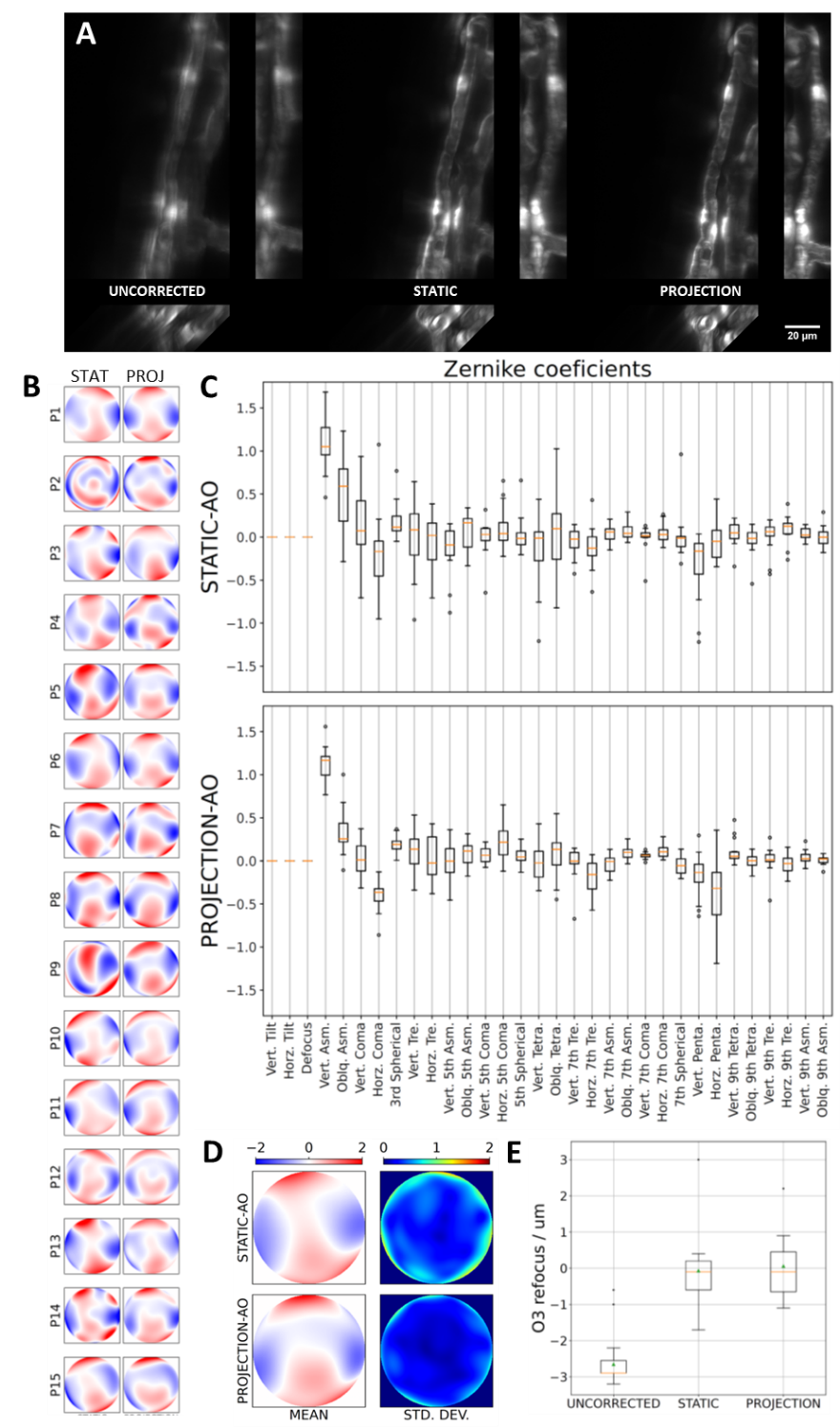

**Supplementary Fig. 4.** Zebrafish imaging within a tube made of fluorinated ethylene propylene (FEP). **A.** Example images of Zebrafish vasculature with system correction, static AO and projection AO. **B.** wavefronts obtained on 15 different positions with both static and projection AO. **C.** Zernike Coefficients corresponding to **B**. **D.** Mean and standard deviations of the wavefronts over all positions, measured using both methods. **E.** Focus correction required for all AO conditions.
